# Down regulation of *Hmgcr* in response to Influenza A infection is independent of the IFN response in human cells

**DOI:** 10.1101/650465

**Authors:** Hongjin Lu, Simon Talbot

## Abstract

Previous studies have demonstrated that the product of the Interferon stimulated gene *Ch25h*, 25-Hydroxycholestrol, provides an immediate and rapid mechanism for down-regulating sterol biosynthesis, through the inhibition of *Hmgcr* gene expression and proteolytic degradation of HMGCR protein. Further studies provide evidence that inhibition of the sterol biosynthesis pathway by 25-HC has broad antiviral effects. In this study, Influenza A virus (IAV) replication was inhibited in cells treated with Fluvastatin or where *Hmgcr* expression was inhibited with an siRNA. Treatment of A549 cells with 25-HC however, resulted in a 2-fold enhancement of IAV replication despite the fact that 25-HC promotes the proteolytic degradation of HMGCR. A549 cells infected with IAV revealed a rapid loss of HMGCR protein, a reduction in *Hmgcr* gene expression as well as an increase in the expression of *Ch25h*, that were all independent of the IFN pathway. Infection of both wild-type and *Ch25h*^*−/−*^ murine BMDMs with IAV also revealed a rapid loss of HMGCR abundance indicating that 25-HC independent mechanisms exist for promoting proteolytic degradation of HMGCR. These data for the human A549 cell line contrast with the induction of *Ch25h* and subsequent loss of HMGCR in murine cells that has been shown to be dependent on IFN signalling.

**Importance:** Cholesterol, a lipid, is mainly produced by the liver or obtained from everyday foods. It is an essential element of the structure of cells and is vital for maintaining the normal function of human body. In cells, cholesterol can be oxidised to oxysterols by biological catalysts, known as enzymes. Certain oxysterols have recently emerged as important elements in the immune response to micro-organisms. This project studied a key enzyme, which is a component of the cholesterol metabolism, known as 3-hydroxy-3-methylglutaryl-CoA reductase (HMGCR). A set of experiments were designed and performed to study the behaviour of HMGCR following viral infections. This study lays the foundation for re-building a map of cholesterol metabolism and the immune response to viral infection. It will be useful for understanding the importance of cholesterol metabolism in infection and exploring novel antiviral strategies.

## 1 Introduction

The sterol metabolism pathway is an intrinsic component of the host defence against infection and has been shown that viral infection or the treatment of murine macrophages with type I or II interferon (IFN) down-regulates the whole sterol biosynthesis pathway [1]. In macrophages and dendritic cells (DCs) viral infection or IFN treatment induces the expression of an interferon stimulated gene, *cholesterol 25-hydroxylase* (*Ch25h*), *via Toll*-like receptor (TLR) induced signalling processing [2, 3].

*Ch25h* catalyses the synthesis of 25-hydroxycholesterol (25-HC), which interacts with the insulin-induced gene proteins (INSIGs). INSIGs retain Sterol Regulatory Element Binding Protein 2 (SREBP2) and the cholesterol-sensing protein SREBP Cleavage-Activating Protein (SCAP) in the ER, suppressing the SREBP pathway [4]. Moreover, 25-HC also induces the ubiquitination and proteasomal degradation of HMGCR, blocking sterol biosynthesis, via SREBP-dependent and SREBP-independent mechanisms [5–7]. However, current data indicate that 25-HC does not play a major role in cholesterol homeostasis but instead has immune modulatory functions [3, 8, 9].

25-HC has antiviral activity against several enveloped viruses by blocking viral entry by directly modifying membranes to inhibit membrane fusions [10]. It also has effects against non-enveloped viruses, including human papillomavirus-16 (HPV-16), human rotavirus (HRoV), and human rhinovirus (HRhV) [11]. 25-HC has also been shown to inhibit differentiation of human monocytes [12], reduce IgA production [13] and alter inflammatory responses [9, 14]. Thus 25-HC is an important component of the IFN-mediated innate immune response to infection but its various roles in immune defence are not completely understood.

In this study, we investigated the replication of Influenza A virus (IAV) in human lung epithelial A549 cells in response to inhibition of the sterol biosynthesis pathway and further examine the expression of both *Hmgcr* and *Ch25h* in response to IAV infection or IFN stimulation in A549 cells and murine BMDMs.

## 2 Experimental procedures

### 2.1 Chemical reagents

25-HC was purchased from Sigma (H1015, Sigma-Aldrich). Mevastatin was purchased from Sigma (M2537, Sigma-Aldrich). Murine recombinant IFN-γ was purchased from Perbio Science or Life Technology (PMC4033).

### 2.2 Cell culture media

**Medium A**: DMEM supplemented with 5% (v/v) CS plus 0.3 mg/ml L-glu; **Medium B**: DMEM supplemented with 5% (v/v) FBS plus 0.3 mg/ml L-glu; **Medium C**: DMEM supplemented with 10% (v/v) FBS plus 0.3 mg/ml L-glu; **Medium D**: DMEM supplemented with 3% (v/v) lipoprotein deficient serum medium (LPDS) (Sigma-Aldrich, UK) plus 0.3 mg/ml L-glu; **Medium E**: medium D plus 0.01 *µ*M of mevastatin; **Medium F**: DMEM-F12 with glutaMAX (Invitrogen, UK) supplemented with 3% (v/v) LPDS, 10% L929 containing colony-stimulating factor 1 (Csf1) and 1.3% (w/v) Penicillin/ 0.6% (w/v) streptomycin; **Medium G**: DMEM-F12 with glutaMAX supplemented with 3% (v/v) LPDS, 0.01 *µ*M of mevastatin, 10% (v/v) L929 containing Csf1 and 1.3% (w/v) Penicillin/ 0.6% (w/v) streptomycin; **Medium H**: DMEM-F12 with glutaMAX supplemented with 10% (v/v) FBS, 10% (v/v) L929 containing Csf1 and 1.3% (w/v) Penicillin/ 0.6% (w/v) streptomycin; **Medium I**: DMEM supplemented with 0.3 mg/ml L-glu.

### 2.3 Mice

C57BL/6 mice were housed in the specific pathogen-free animal facility at the University of Edinburgh. *Ch25h*^*−/−*^ (B6.129S6-Ch25h^tm1Rus^/J) mice were purchased from Charles River (Margate, United Kingdom) and housed in the specific pathogen-free animal facility at the University of Edinburgh [15].

### 2.4 Cell Propagation and Culture

The NIH/3T3 fibroblast cell line (ATCC CRL1658) was obtained from American Type Culture Collection (ATCC) and grown in medium A. A549 cells were grown in medium B. MDCK (Madin-Darby canine kidney) cells were grown in medium C. The A549/PIV5-V cell line was kindly provided by Professor Richard Edward Randall (University of St Andrews), which was grown in medium B. Wild-type BMDMs were derived from the femur and tibia isolated from C57BL/6 mice as previously described [1, 3]. *Ch25h*^*−/−*^ BMDMs were derived from the femur and tibia isolated from B6.129S6*Ch25h*^tm1Rus^/J mice and grown in medium H. All procedures were carried out under project and personal licences approved by the Secretary of State for the Home Office, under the United Kingdoms 1986 Animals (Scientific Procedures) Act and the Local Ethical Review Committee at Edinburgh University. All cultures are routinely tested for mycoplasma and endotoxin levels.

### 2.5 Viruses

Murine cytomegalovirus (MCMV) has been previously described [1]. Influenza A/WSN/1933(H1N1) virus was propagated and titred in MDCK cells.

### 2.6 qRT-PCR analysis

QIAGEN Rneasy Plus Mini Kit (QIAGEN, Germany) was used to purify total RNA. qScript One-Step Fast qRT-PCR kit, Low ROX (95081-100, Quanta BioSciences, USA) and Bioline SensiFAST SYBR Lo-ROX One-Step Kit (BIO-74005, Bioline, USA) were used for the qRT-PCR measurement. qRT-PCR was undertaken according to manufacturer’s instructions. All Taqman probe/primer systems and primers used here are listed in Appendix Table 1 and Table 2.

### 2.7 qRT-PCR data analysis

The MxPro qPCR software (Stratagene, USA) can be used to analyse the qRT-PCR data. Alternatively, Ct values can be exported from the software and Livak method (also known as Delta Delta CT method) [16] can be used to analyse the qRT-PCR results, which is shown below:

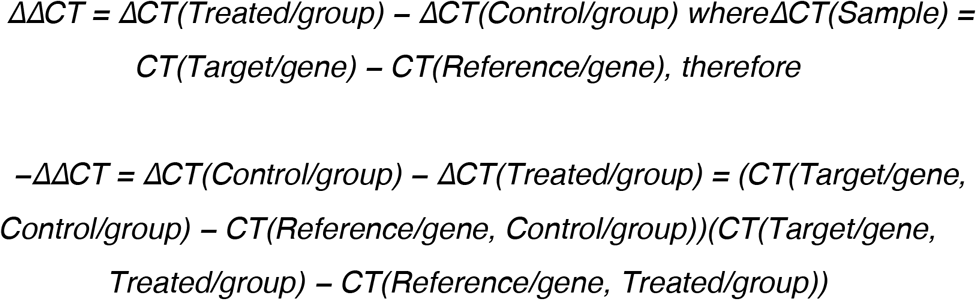

Thereafter, the ratio of normalised target gene in treated groups relative to that in control groups can be calculated using:

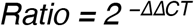

### 2.8 siRNA transfection

DharmaFECT 1 Transfection Reagent (Thermo Scientific, UK) was used for siRNA transfection. Experiments were performed according to manufacturer’s instructions. siRNAs are listed in Appendix Table 3.

### 2.9 Western blot analysis

Cells were seeded in 24-well plates. After treatments, cells were lysed directly in wells by adding 100 *µ*l of RIPA lysis buffer (9806, Cell Signaling) supplemented with protease cocktail (cOmplete Protease Inhibitor Cocktail Tablets, Roche) and 1 mM Phenylmethanesulfonyl fluoride (Sigma, UK). Plates were incubated on ice for 20 minutes. Protein concentrations of whole-cell lysate fractions were then determined using the BCA Protein Assay Reagent kit (Thermo Scientific) according to manufacturer’s instructions.

Prior to SDS-PAGE, 20 *µ*g to 50 *µ*g of protein for each sample was mixed with 2X Laemmli sample buffer (S3401, Sigma) and heated at 50°C for 10 minutes. Thereafter, mixtures were subjected to 8% SDS-PAGE with 6 *µ*l of molecular weight marker (SeeBlue Plus2 Pre-Stained Standard, Invitrogen), after which proteins were transferred to nitrocellulose membranes (Amersham Hybond ECL 0.45 *µ*m Nitrocellulose Membrane, GE Healthcare Life Sciences) or PVDF membranes (Amersham Hybond P 0.45 *µ*m PVDF Membrane, GE Healthcare Life Sciences). The membranes were blocked with 5% skimmed milk (Sigma, UK), probed with specific primary antibodies overnight at 4°C, washed with PBST, incubated with HRP-conjugated secondary antibodies at room temperature for 1 hour. The membranes were then re-washed with PBST and bands were visualized using X-ray films (Amersham Hyperfilm ECL, UK) or Odyssey Fc Imager (LI-COR Biosciences, UK) (kindly provided by Professor Seth Grant, University of Edinburgh). Image Studio Lite (Li-COR Biosciences, UK) was used to analyze the bands. Antibodies used here are listed in Appendix Table 4.

#### 2.9.1 Preparation of HMGCR monoclonal antibody

A9 hybridoma was purchased from ATCC (CRL1811). Cells were recovered immediately and grown in medium C. In order to prepare HMGCR monoclonal antibody, cells were centrifuged and re-suspended in DMEM supplemented with 2.5% (v/v) FBS plus 0.3 mg/ml L-glu or Gibco Hybridoma-SFM (12045076, Life Technologies, UK). 20 ml of 2×10^5^ cells/ml were transferred to a 75 ml flask. Cells were then kept growing in Hybridoma-SFM until more than 95% cells were dead. The supernatant was collected, aliquoted and frozen at −20°C. Western blot was then performed to determine the optimal dilution factor of the HMGCR monoclonal (A9) antibody.

## 3 Results

### 3.1 Replication of influenza A virus in response to HMGCR expression levels

Previous studies indicate that during virus infection 25-HC provides an immediate and rapid mechanism for down-regulating sterol biosynthesis, which involves a negative feedback regulation of the sterol biosynthesis pathway and the proteolytic degradation of HMGCR via the IFN signalling pathway [3, 7]. In addition to the critical role in manipulating sterol biosynthesis, HMGCR is also a target for 25-HC associated IFN-mediated host defence against viral infection. However, it remains unknown whether HMGCR is the only target of 25-HC associated IFN-mediated innate immune response and whether this immune response broadly exists in host defence against infection. Experiments were performed to investigate whether 25-HC inhibits Influenza A virus (IAV) infection through the IFN-mediated innate immune response. Figure 1A and B show that the treatment of A549 cells (human lung epithelial cells) with 2.5 *µ*M 25-HC enhanced the replication of IAV, in contrast to previously published data showing a reduction of replication of IAV in MDCK cells (canine kidney cell line) following 25HC treatment [1]. However, when A549 cells were treated with Fluvastatin to inhibit the action of HMGCR, a dose dependent reduction in IAV replication of up to 50% was observed (Figure 1B). The effect of siRNA-mediated knockdown of *Hmgcr* on IAV replication in A549 cells (Figure 1C) resulted in 15% reduction in IAV titres. These results show that although both pharmacological and siRNA mediated knockdown of HMGCR have an antiviral effect, in contrast, 25-HC promotes IAV replication in A549 cells.

**Figure 1:**
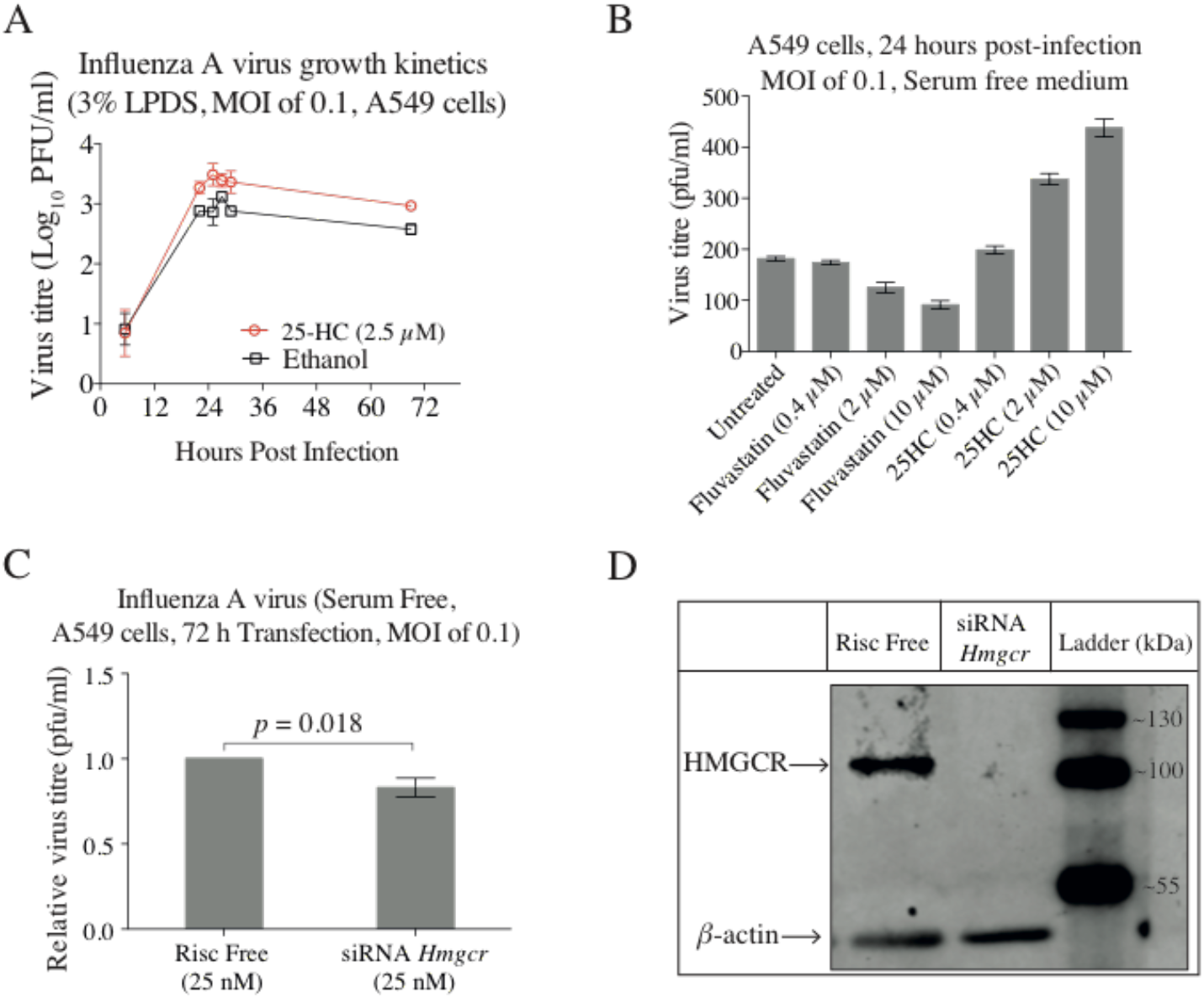
Viral replication in *Hmgcr*-deficient cells. **A.** Effects of 25-HC on IAV growth kinetics. Ethanol was used as a negative control. **B.** A549 cells were infected with IAV at a MOI of 0.1 in medium I for one hour. Virus was then removed and cells were treated with 0.4 *µ*M, 2 *µ*M and 10 *µ*M of fluvastatin or 25-HC in medium I, respectively. Supernatant was collected at 24-hour post-infection. A plaque assay was performed to determine virus titres. Data are ± SEM (n = 3 biological replicates). **C.** A549 cells were transfected with siRNA *Hmgcr* for 72 hours in medium E. One group of cells were then harvested for western blot. Another group was infected with IAV at a MOI of 0.1 in medium I for another 24 hours. A plaque assay was performed to determine virus titres at 24-hour post-infection. Risc free acts as a negative control. Data are ± SEM (n = 8 technical replicates). **D.** Western blot was carried out to validate the HMGCR protein levels. β-actin was used as a loading control. C-1 HMGCR antibody was used for protein detection. IAV infection leads to a reduction in *Hmgcr* gene expression and HMGCR protein abundance.

### 3.2 IAV infection leads to a reduction in *Hmgcr* gene expression and HMGCR protein abundance

To investigate whether IAV infection down-regulates HMGCR gene expression and endogenous protein levels, A549 cells were treated and infected as indicated in Figure 2. A low concentration of mevastatin (0.01 *µ*M) was used to allow for an increased dynamic range for measuring levels of HMGCR [7]. Figure 2 shows that whilst a low concentration of mevastatin strongly induced the biosynthesis of HMGCR, viral infection resulted in dramatic reduction in HMGCR protein levels (Figure 2A and B). In contrast, *Hmgcr* gene expression levels were moderately increased by mevastatin treatment in both uninfected and infected groups. A dramatic reduction in *Hmgcr* gene expression following IAV infection was observed at 24-hour post-infection (Figure 2C). In addition, viral infection induced *Ch25h* gene expression, which reached a peak at 8-hour post-infection and then decreased (Figure 2D). Taken together, IAV infection leads to a reduction in *Hmgcr* gene expression at late times, whereas the level of HMGCR protein quickly and dramatically dropped following infection. These data suggest that at early times the reduction in HMGCR abundance was mainly due to protein degradation rather than transcriptional down-regulation.

**Figure 2.**
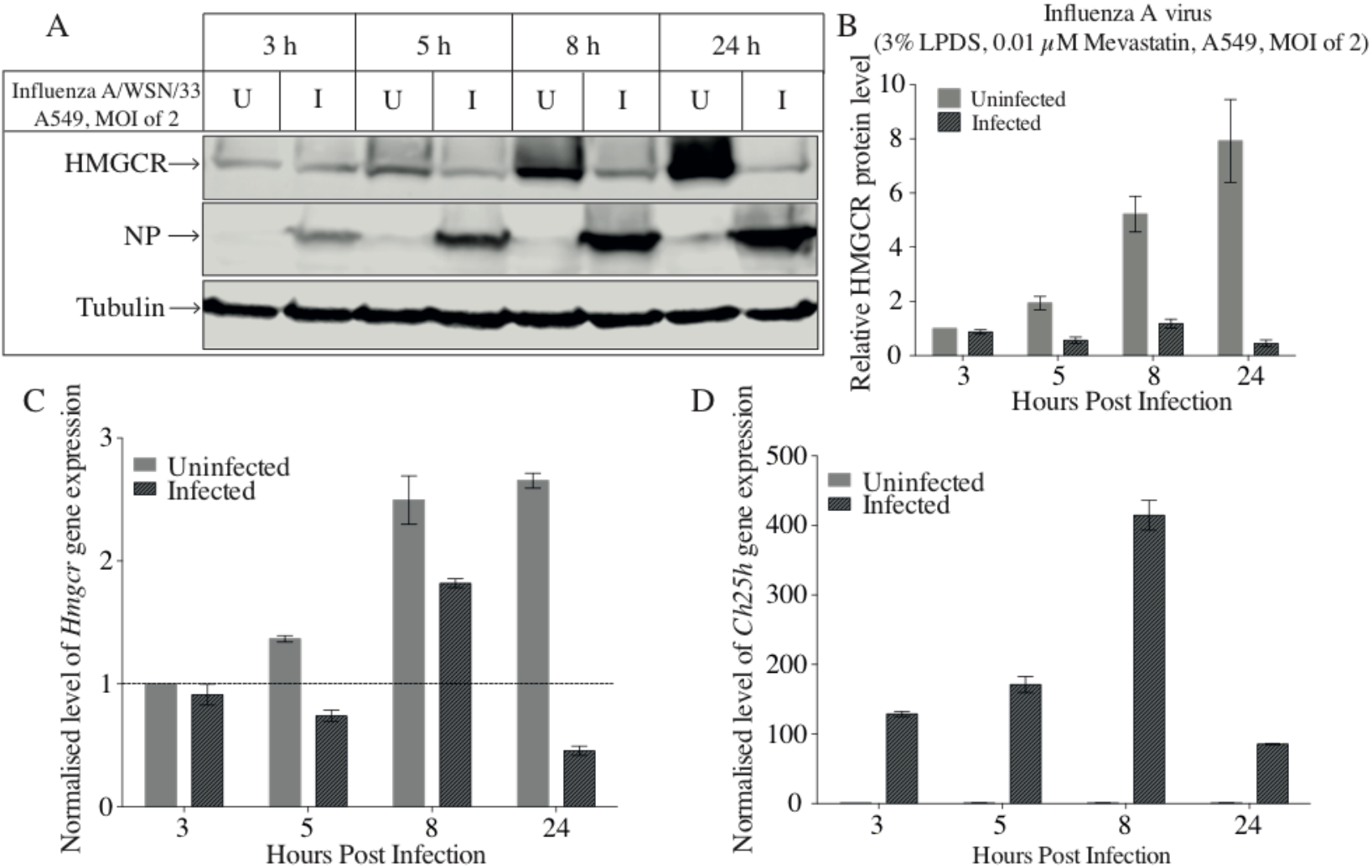
Time course study of HMGCR protein levels with IAV infection in A549 cells. A549 cells were pre-treated with medium E overnight. The next day, cells were infected with IAV at a MOI of 2 in medium E. Cells were harvested at indicated time points (post-treatment). **A.** Western blot was performed to determine HMGCR and NP protein levels. C-1 HMGCR antibody was used. Tubulin was used as the loading control. **B.** Intensity values of HMGCR to tubulin were calculated. **C.** qRT-PCR was used to quantify *Hmgcr* gene expression levels. **D.** Quantification of *Ch25h* gene expression levels. Bars present ± SEM (n = 3 technical replicates). U: uninfected; I: infected.

### 3.3 Is the reduction of HMGCR mediated by newly synthesised 25-HC following viral infection?

Wild-type and *Ch25h*^*−/−*^ murine BMDMs were infected with IAV and HMGCR protein levels were measured at indicated time points. Cells were pre-treated with lipoprotein-deficient medium to increase the level of HMGCR protein. The basal level of HMGCR in *Ch25h*^*−/−*^ BMDMs is high enough to be detected under mevastatin-free conditions.

In wild-type BMDMs IAV infection greatly decreased HMGCR protein levels beginning at 3 hours post-infection (Figure 3A). In *Ch25h*^*−/−*^ BMDMs the decrease of HMGCR was first observed at 8 hours post-infection (Figure 3B). The decrease of HMGCR protein level is delayed in *Ch25h*^*−/−*^ BMDMs, suggesting that the newly synthesised 25-HC is most likely responsible for the reduction of HMGCR after infection at early time points (< 5 hours). However, the decrease after 8 hours post-infection cannot be caused by the action of 25-HC, since this was also observed in *Ch25h*^*−/−*^ BMDMs. These data support a role for both 25HC dependent and independent mechanisms for the loss of HMGCR abundance in response to IAV infection in human cells.

**Figure 3.**
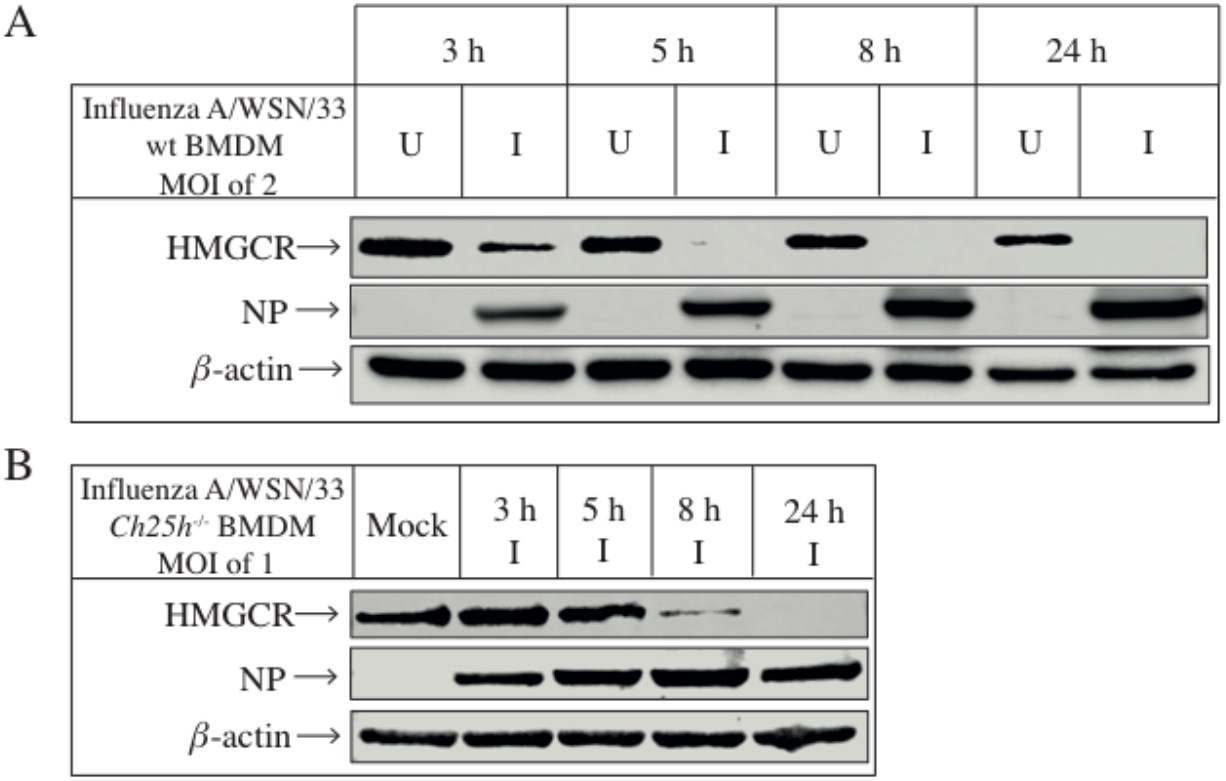
Time course study of HMGCR protein levels with IAV infection in wild-type and *Ch25h*^*−/−*^ BMDMs. **A.** Wild-type BMDMs were pre-treated with medium G overnight and then infected with influenza virus at a MOI of 2 in medium G. Cells were harvested at indicated time points (post-treatment). Western blot was performed to determine HMGCR and NP protein levels in wild-type BMDMs. β-actin is the loading control. C-1 HMGCR antibody was used. **B.** *Ch25h*^*−/−*^ BMDMs were infected with influenza virus at a MOI of 1 in medium F. Samples were collected for western blot. C-1 HMGCR antibody was utilized for HMGCR detection. β-actin is the loading control. U: uninfected; I: infected.

### 3.4 Is the degradation of HMGCR induced by IAV infection dependent on the IFN pathway?

The above experiments indicate that 25-HC is not exclusively responsible for the down-regulation of HMGCR during viral infection. Further experiments were then performed to study whether the degradation of HMGCR induced by IAV infection is dependent on the IFN pathway in human cells. A549 cells were treated with IFN-α or IFN-γ. Figure 4A and C show that, in contrast to virus infection and 25-HC treatments which dramatically reduced HMGCR protein levels, treatment with IFN-α or IFN-γ only modestly reduced HMGCR protein levels. This contrasts with the treatment of murine BMDMs with IFN-γ that leads to a dramatic reduction in HMGCR levels (Figure 4F). In order to validate this result, an A549/PIV5-V cell line, whose IFN signalling pathway is blocked by targeting the STAT1 for proteasome-mediated degradation [17], was infected with IAV (Figure 4E). HMGCR protein levels greatly decreased in both A549 cells and A549/PIV5-V cells infected with IAV, suggesting that at 24-hour post-infection the STAT-mediated IFN response is not the main reason for the reduction of HMGCR during IAV infection.

**Figure 4.**
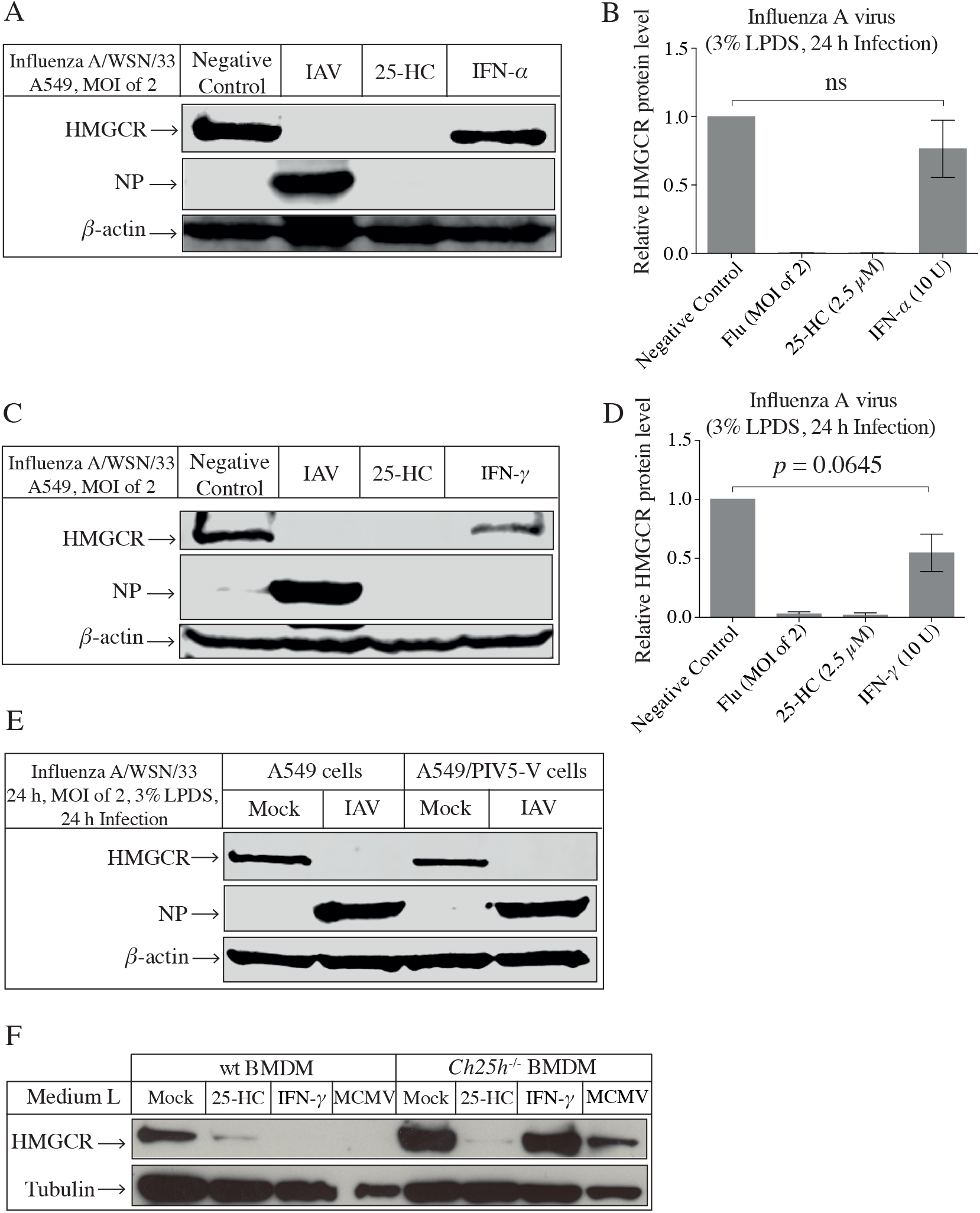
The endogenous HMGCR protein levels in A549 or A549/PIV5-V cells with treatments. **A.** A549 cells were treated with IAV (MOI of 2), 25-HC (2.5 *µ*M) and IFN-α (10 U) in medium D, respectively. After 24-hour post-treatment, cells were harvested for western blot. A9 HMGCR antibody was used. β-actin is the loading control. **B.** Intensity values of HMGCR to β-actin. Bars present ± SEM (n = 3 technical replicates). An unpaired Student’s *t* test with Welch’s correction was performed. **C.** A549 cells were treated with IAV (MOI of 2), 25-HC (2.5 *µ*M) and IFN-γ (10 U) in medium D, respectively. After 24-hour post-treatment, cells were harvested for western blot. A9 HMGCR antibody was used. β-actin is the loading control. **D.** Intensity values of HMGCR to β-actin. Bars present ± SEM (n = 4 technical replicates). An unpaired Student’s *t* test with Welch’s correction was performed. **E.** Cells were infected with IAV at a MOI of 2 in medium D and harvested for western blot at 24-hour post-infection. A9 HMGCR antibody was used. β-actin is the loading control. **F**. Wild-type and *Ch25h*^*−/−*^ BMDMs were treated with 2.5 *µ*M of 25-HC, 10 U of IFN-γ or infected with MCMV at a MOI of 2 in medium G, respectively. Cells were harvested at 24-hour post-treatment for the determination of HMGCR protein levels. C-1 HMGCR antibody was used. Tubulin was used as a loading control.

### 3.5 How is *Hmgcr* gene expression affected by IAV infection or IFN treatment?

The expression of *Hmgcr*, *Ch25h* and the Interferon stimulated gene *Mx1* was examined after either treatment with IFN-α, IFN-γ or infection with IAV in A549 or A549/PIV5-V cells. Treatment of A549 cells with IFN-α had no effect on the expression of *Hmgcr* or *Ch25h*, whereas there was a modest increase of expression of *Hmgcr* following IFN-γ treatment (Figure 5A and C). Infection of A549 cells with IAV on the other hand, led to a sharp increase in *Ch25h* expression and a decrease in *Hmgcr* expression (Figure 5A and C). Similar changes to the levels of expression of *Ch25h*, an *Hmgcr* in response to IAV infection were also observed in A549/PIV5-V cells that are unable to respond to IFN stimulation (Figure 5B and D). These results support the finding that IAV infection of A549 cells leads to a reduction in *Hmgcr* expression and HMGCR abundance through an IFN-independent pathway.

**Figure 5.**
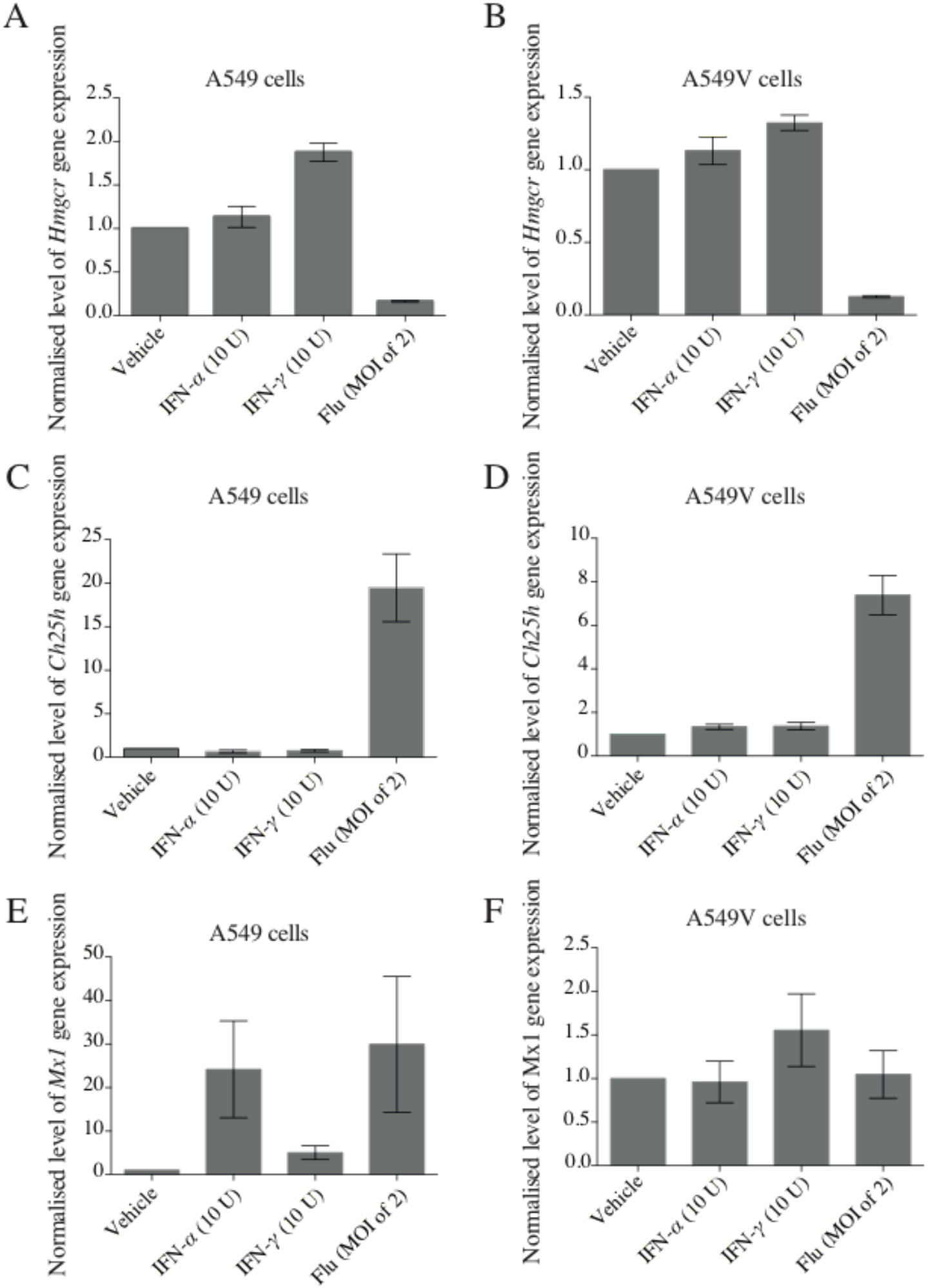
The expression levels of genes in A549 cells with treatments. A549 cells and A549/PIV5-V cells were treated with IFN-α (10 U), IFN-γ (10 U) and IAV (MOI of 2) in medium D, respectively. Cells were harvested at indicated time points and qRT-PCR was then performed to determine the expression levels of various genes. **A.** *Hmgcr* gene expression levels in A549 cells at 24 hours post-treatment. **B.** *Hmgcr* gene expression levels in A549/PIV5-V cells at 24 hours post-treatment. **C.** *Ch25h* gene expression levels in A549 cells at 24 hours post-treatment. **D.** *Ch25h* gene expression levels in A549/PIV5-V cells at 24 hours post-treatment. **E.** *Mx1* gene expression levels in A549 cells at 5 hours post-treatment. **F.** *Mx1* gene expression levels in A549/PIV5-V cells at 5 hours post-treatment. Target genes were normalised by multiples internal control genes, including *Gapdh, Hmbs* and *Rpl13a* [18]. Bars present ± SEM (n = 4 technical replicates). An unpaired Student’s *t* test with Welch’s correction was performed.

## 4 Discussion

25-HC shows antiviral effects on a wide range of viruses by inhibiting different steps during viral infection, such as blocking viral entry and inhibiting the viral replicative cycle [3, 19–21]. In this study we show that 25-HC promotes the replication of IAV in a human lung epithelial cell line (A549) (Figure 1) in a dose dependent manner (> 1 *µ*M). These results conflict with previously published data showing that IAV replication is inhibited by 25-HC in other cells such as MDCK (Canine kidney) and Murine lung epithelial cells [9]. Specifically targeting the sterol biosynthesis pathway through the use of statins or siRNAs directed against HMGCR proved to be antiviral, suggesting that the pro-viral effects of 25-HC must be due to the influence on other pathways or processes. For example, 25-HC has a role in altering cell inflammatory responses [9] and affecting inflammasome activity [14]. It could be speculated that 25-HC also influences the expression of viral restriction factors.

The RNAi mediated knockdown of HMGCR protein levels and pharmaceutical inhibition of HMGCR activity down-regulates viral activity (Figure 1) and have the potential to protect hosts from viral infection [1, 22–25]. Moreover, the inhibition of HMGCR increases the expression of interferon-responsive genes [26]. It has been reported that the changes in the metabolic flux of the sterol biosynthesis pathway elicit a type I IFN response mediated by STING (stimulator of interferon genes) [27]. In Murine models at least, the IFN response down-regulates sterol biosynthesis via increasing the biosynthesis of 25-HC, which also enhances the efflux of sterols, and down-regulation of the sterol biosynthesis pathway results in the up-regulation of the type I IFN response [27]. A recent paper shows that the anti-dengue virus properties of statins may associate with the down-regulation of the expression of cellular immune and pro-inflammatory response related genes, which is independent of cholesterol levels [28]. These studies suggest that HMGCR plays a role in the hosts’ response to viral infection. Based on these findings, we have further investigated the relationship between HMGCR and the expression of 25-HC in response to viral infection. Figure 2 and Figure 3 show that IAV infection down-regulated HMGCR protein levels. However, at late times (> 8 hours) the down-regulation of HMGCR observed in *Ch25h*^*−/−*^ BMDMs suggests that this reduction is mediated through 25-HC independent mechanisms (Figure 3). Moreover, Figure 4 shows that in human A549 cells, HMGCR down-regulation is IFN independent, as the reduction of HMGCR during IAV infection occurred in STAT1-deficient cells (Figure 4). Furthermore, we show that IAV infection stimulated the gene expression of *Ch25h* and reduced HMGCR protein level at 24 hours post-infection, in contrast to IFN treatment (Figure 4 and Figure 5). These results indicate that the decrease of HMGCR post-infection is probably mediated by *de novo* synthesised 25-HC, but the biosynthesis of 25-HC is independent of the IFN signalling pathway in human cells. In another study it was demonstrated that 25-HC inhibits the genome replication of Hepatitis C virus (HCV) by suppression of the SREBP pathway in humans. However, the up-regulation of CH25H in human cells is not induced by the IFN response; instead it is induced by intrinsic innate immune responses [29]. The data presented here are thus in good agreement with findings that the *de novo* synthesis of 25-HC during IAV infection in A549 cells and down regulation of HMGCR is independent of the IFN-signalling pathway.

Results presented in this study indicate that, during IAV infection, alternative IFN and/or 25-HC independent mechanisms or pathways also contribute to the down-regulation of HMGCR during infection. This study provides further understanding to the role of 25-HC in innate immunity and the crosstalk between the sterol metabolism and innate immune response against viral infection. Further studies are required to understand the sterol metabolism-IFN signalling pathway circuit in innate immunity and explore other IFN-independent pathways that contribute to the down-regulation of sterol metabolism in viral infection.

## Acknowledgements

HL was supported by the China Scholarships Council/University of Edinburgh Scholarships. We would like to thank Professor Peter Ghazal for supervision. We are grateful to Professor David Russell (UT Southwestern, Dallas, Texas, USA) for the supply of the *Ch25h*^*−/−*^ mice. We would like to thank Marie Craigon and Alan Ross for technical support.

## Appendix

**Table 1.**
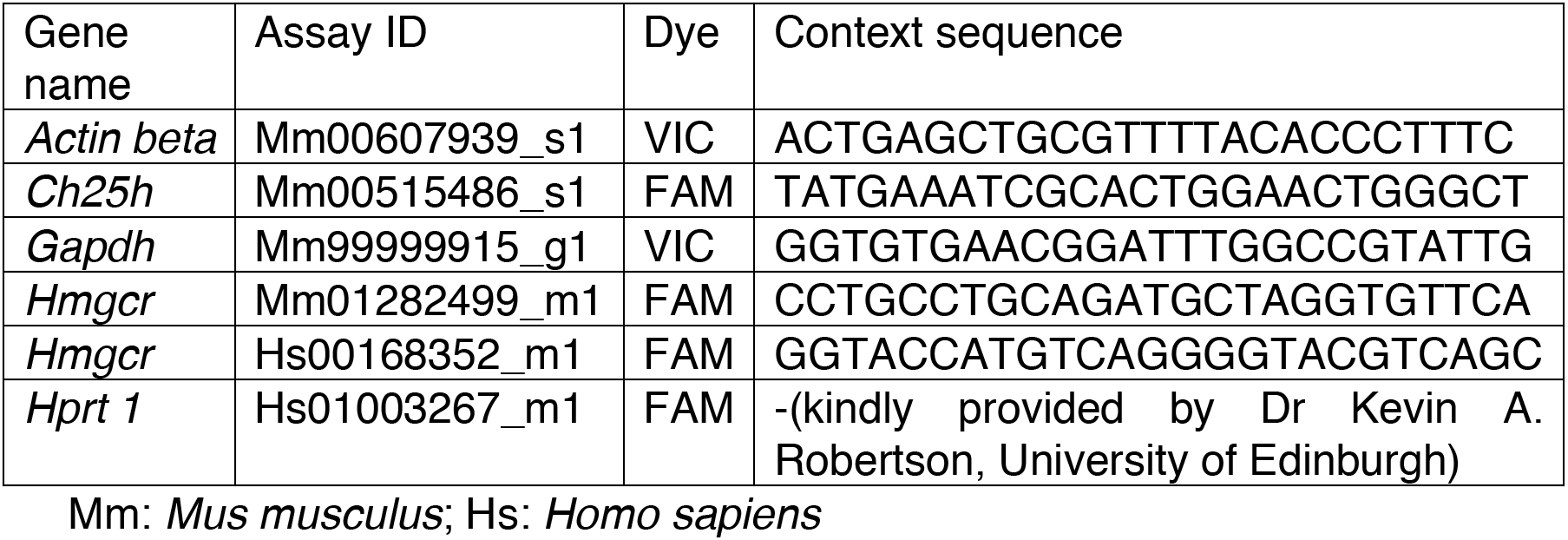
Information for Taqman probe/primer systems

**Table 2.**
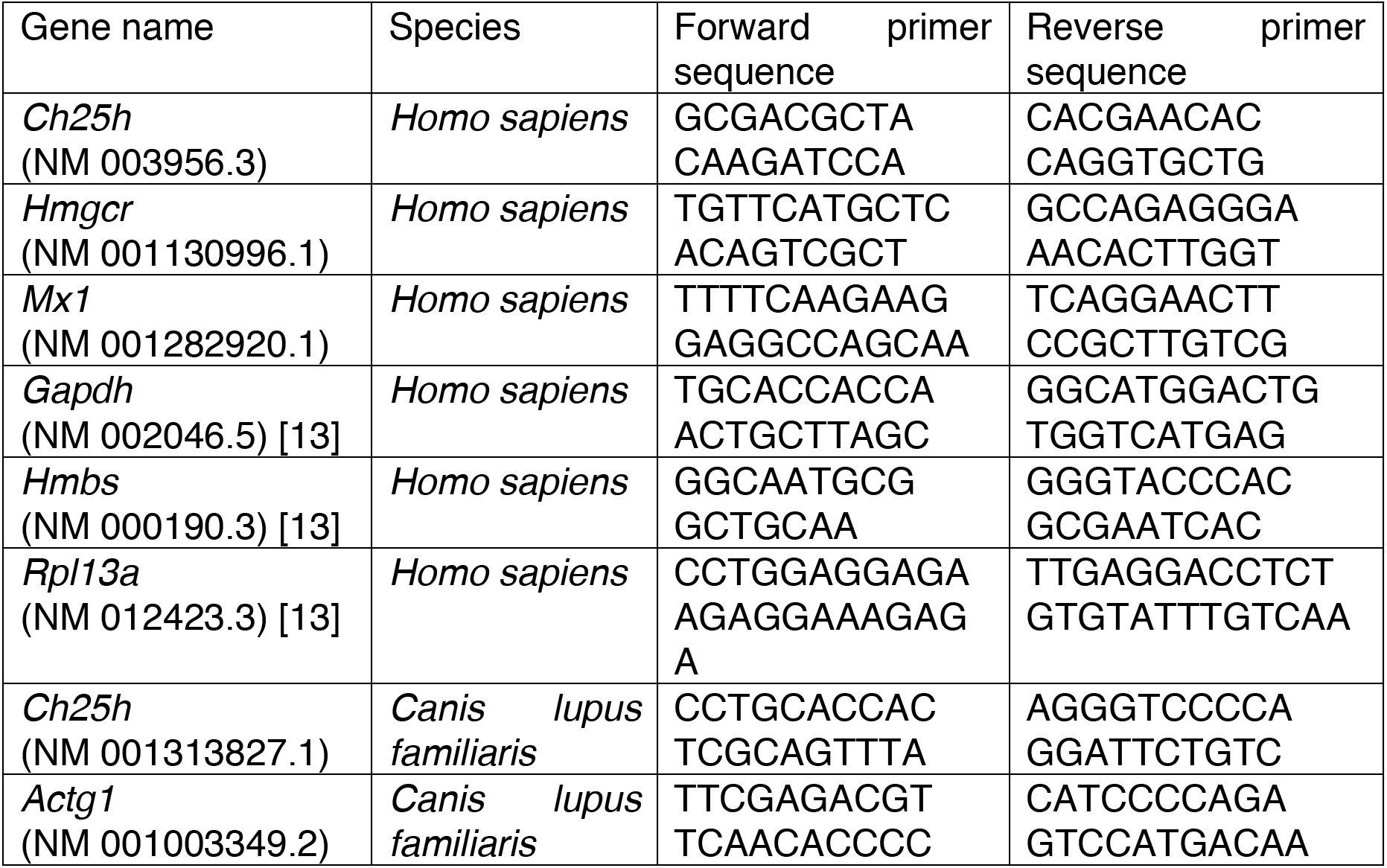
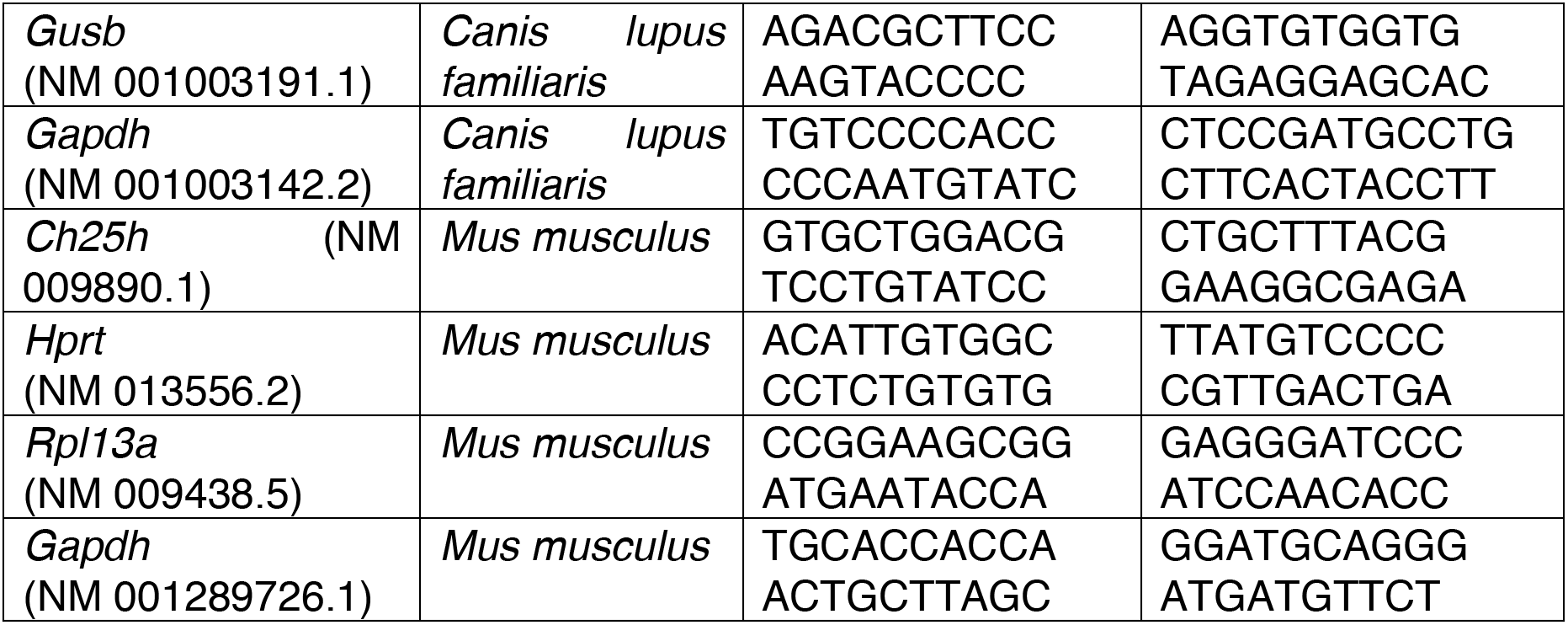
Primers information

**Table 3.** siRNA information

| Name | Catalogue Number | Species |
| --- | --- | --- |
| siRNA HMGCR | LU-009811-00-0002 (Thermo Scientific, UK) | *Homo sapiens* |
| siRNA HMGCR | LU-040731-01-0002 (Thermo Scientific, UK) | *Mus musculus* |
| Risc free | D-001600-01 (Thermo Scientific, UK) | *Mus musculus* |

**Table 4.**
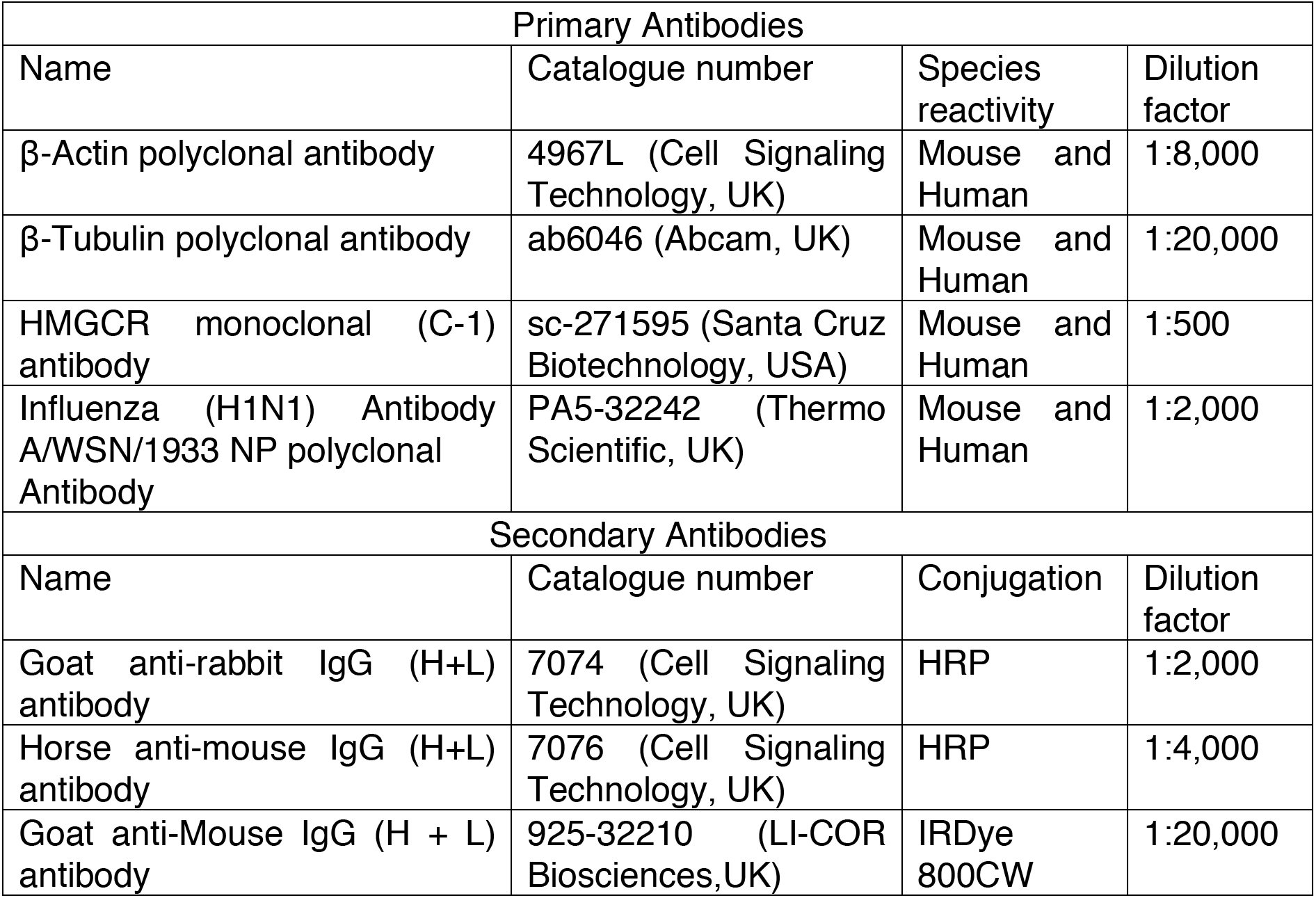
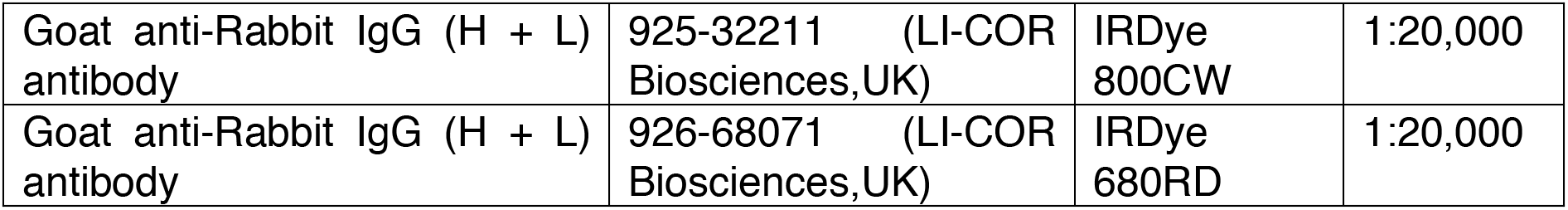
Antibodies for western blot

